# Insights into telomerase action from high-throughput sequencing of *S. pombe* telomeres

**DOI:** 10.1101/047837

**Authors:** Henrietta W. Bennett, Na Liu, Yan Hu, Megan C. King

## Abstract

We have developed a high-throughput sequencing approach that enables us to determine terminal telomere sequences from tens of thousands of individual *Schizosaccharomyces pombe* telomeres. This method provides unprecedented coverage of telomeric sequence complexity in fission yeast. *S. pombe* telomeres are composed of modular degenerate repeats that can be explained by variation in usage of the TER1 RNA template during reverse transcription. Taking advantage of this deep sequencing approach, we find that “like” repeat modules are highly correlated within individual telomeres. Moreover, repeat module preference varies with telomere length, suggesting that existing repeats promote the incorporation of like repeats and/or that specific conformations of the telomerase holoenzyme efficiently and/or processively add repeats of like nature. After the loss of telomerase activity, this sequencing and analysis pipeline defines a population of telomeres with altered sequence content. This approach should be adaptable to study telomeric repeats in other organisms and can provide new insights into telomere sequence content at high density.

## Introduction

The ends of linear chromosomes are protected from degradation and end-to-end fusion by nucleoprotein structures called telomeres. While the 5’ ends of the chromosome can be replicated in their entirety by leading strand synthesis, the inability of the lagging strand synthesis machinery to replicate the very 3’ ends leads to progressive chromosome shortening (1,2). Nucleolytic degradation further contributes to loss of telomeric sequence (3-6). Telomere shortening can be counteracted by telomerase – a ribonucleoprotein complex with reverse transcriptase activity that catalyzes addition of nucleotides to the 3’ end of the chromosome (7). The nucleotide sequence added to the 3’ end is specified by the RNA component of telomerase, leading to telomere ends that are composed of degenerate repeats (8). Proteins that recognize both double-stranded and single-stranded telomeric repeats help to cap telomeres by directly suppressing DNA repair activities (9) or by driving formation of a “T-loop” that effectively sequesters the 3’ ssDNA overhang in some organisms (10). In addition, these telomere binding proteins control recruitment of telomerase, which acts predominantly during late S/G2 phase of the cell cycle when telomeres are replicated (11).

The fission yeast telomerase holoenzyme is composed of three core components: the catalytic component Trt1, the RNA template TER1, and the accessory protein Est1, and is associated with additional factors, such as the Lsm2-8 proteins (12-16). The TER1 template determines the sequence of the telomeric repeat with the most common repeat in *S. pombe* being GGTTACA (13,16); significant degeneracy in these telomere repeat sequences in *S. pombe* (17) suggests that the coordination between the reverse transcriptase activity and the template RNA structure could underlie variability in repeat usage. Although this relationship is not completely understood, the position and base pairing between the 3’ end of the chromosome (effectively the “primer”) and the TER1 RNA, the extent of “G-stutter” that the enzyme undergoes, and the integrity of the template boundary that defines the end of the repeat unit could all contribute to the observed sequence heterogeneity (16,18-20). Heterogeneity in telomere sequence appears common in yeast models (21,22); whether this heterogeneity is biologically important remains an open question.

The inherent repetitive nature of telomeres and the rapid change in telomere content that occurs during cell culturing present challenges to employing high-throughput sequencing methods to interrogate the complexity and regulation of telomere sequence. Indeed, the lack of a reference sequence and inability to uniquely assign repeat-containing reads has thus far prevented assembly of true chromosome ends. Here, we circumvent this challenge by developing and employing a method to sequence individual *S. pombe* telomeres in their entirety. In addition to improving the precision of measuring telomere length within a cell population, the depth of sequencing afforded by this approach has revealed patterns of telomere repeat content within individual telomeres. Further, we found a previously unappreciated connection between the prevalence of particular repeat sequences and telomere length. Moreover, we demonstrate that this approach will be useful for investigating how perturbations to telomere maintenance influence telomere sequence. We anticipate that this method can be adapted to sequence telomeres in a wide variety of organisms (up to approximately 5 kilobase pairs in length) and may be particularly useful for assessing how telomere dysfunction influences telomeric sequence.

## Materials and Methods

### *S. pombe* strain generation and culture conditions

All experiments were carried out on a wild type *S. pombe* strain (h+, ura4-D18, leu1-32, ade6-M216; MKSP200) except for Figure 6 (the *trt1Δ* cells; genotype: h?, leu1-32, ade6?, trt1::KanR; MKSP1424; spores were cultured after dissection from a diploid heterozygous for trt1/trt1::KanR). Cell culturing was carried out as described (23). *S. pombe* were grown at 30°C.

**Figure 1.**
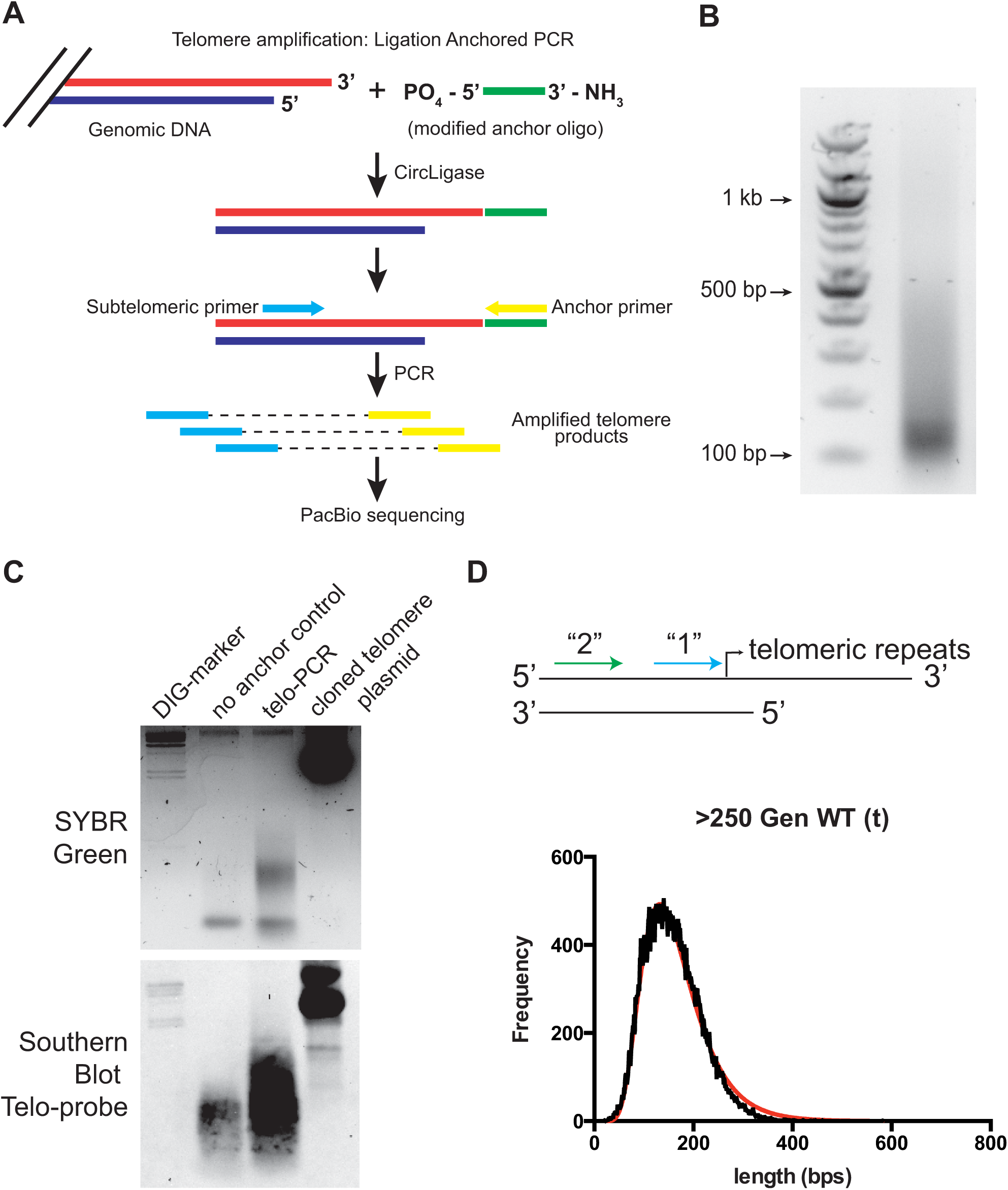
Method for high-throughput telomere sequencing in *S. pombe.* (A) Schematic of the protocol used to amplify the telomeric repeats. An anchor primer is ligated onto chromosome ends using CircLigase followed by PCR with a subtelomeric primer and the reverse complement of the anchor sequence. The products are purified to remove free primers using a Microcon concentrator and used to generate circular libraries for PacBio sequencing. (B) Example of purified telomere PCR products for WT cultures. Genomic DNA was subjected to the procedures outlined in (A) using subtelomeric primer “1”. (C) An example of telo-PCR reactions comparing a “no anchor” control with the same sample in which ligation of the anchor to chromosome ends was first carried out. A plasmid containing a cloned telomere serves as a control. Agarose gel stained with SYBR Green (top) and Southern blot hybridized with a Digoxigenin telomere probe detected with anti-DIG-AP (bottom). (D) Histogram of telomere length for a population of ~60,000 WT telomere sequences. The cartoon at the top schematizes the binding sites for subtelomeric primer “1” (cyan) and “2” (green).

**Figure 2.**
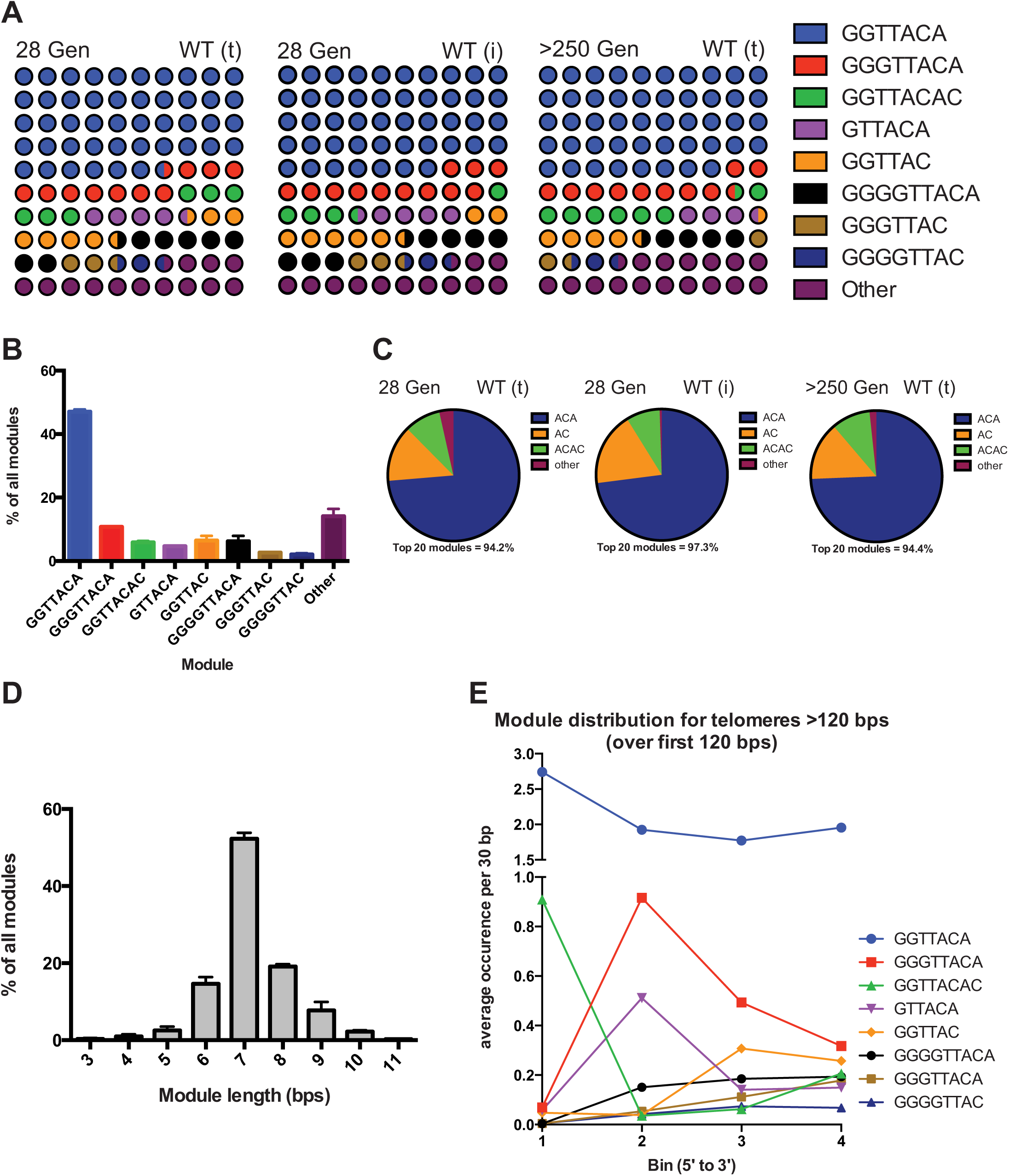
*S. pombe* telomeres are composed of module variants (A) The canonical GGTTACA repeat makes up nearly half of all telomere sequence. Representation of the repeat content of WT telomeres shortly after germination (28 generations) amplified by two different protocols (t) and (i), see text, or after extensive culturing (>250 generations). Each telomere was broken down into its repeat units or “modules”. The eight most common repeats are coded as colors with the rest of the telomere repeats categorized as “other”. Each circle is equivalent to 1%. (B) Plot of the percentage that each module makes up of all modules in the telomere population, plotted as the mean with the standard deviation from the three libraries shown in panel (A). (C) Distribution of the 3’ endings of modules from WT cells amplified shortly after germination (28 generations) using two different amplification protocols (t) and (i), see text, or after extensive culturing (>250 generations). (D) The distribution of module length from the three libraries shown in panel (A), plotted as the means with the standard deviation. (E) Distribution of module usage over the first 120 bps for telomeres >120 bp in length. The 120 bps at the 5’ end of each telomere was split into 4 x 30 bp bins (numbered 1-4 from 5’ to 3’) and the module occurrence was quantified; this data was derived from the >250 generation WT library.

**Figure 3.**
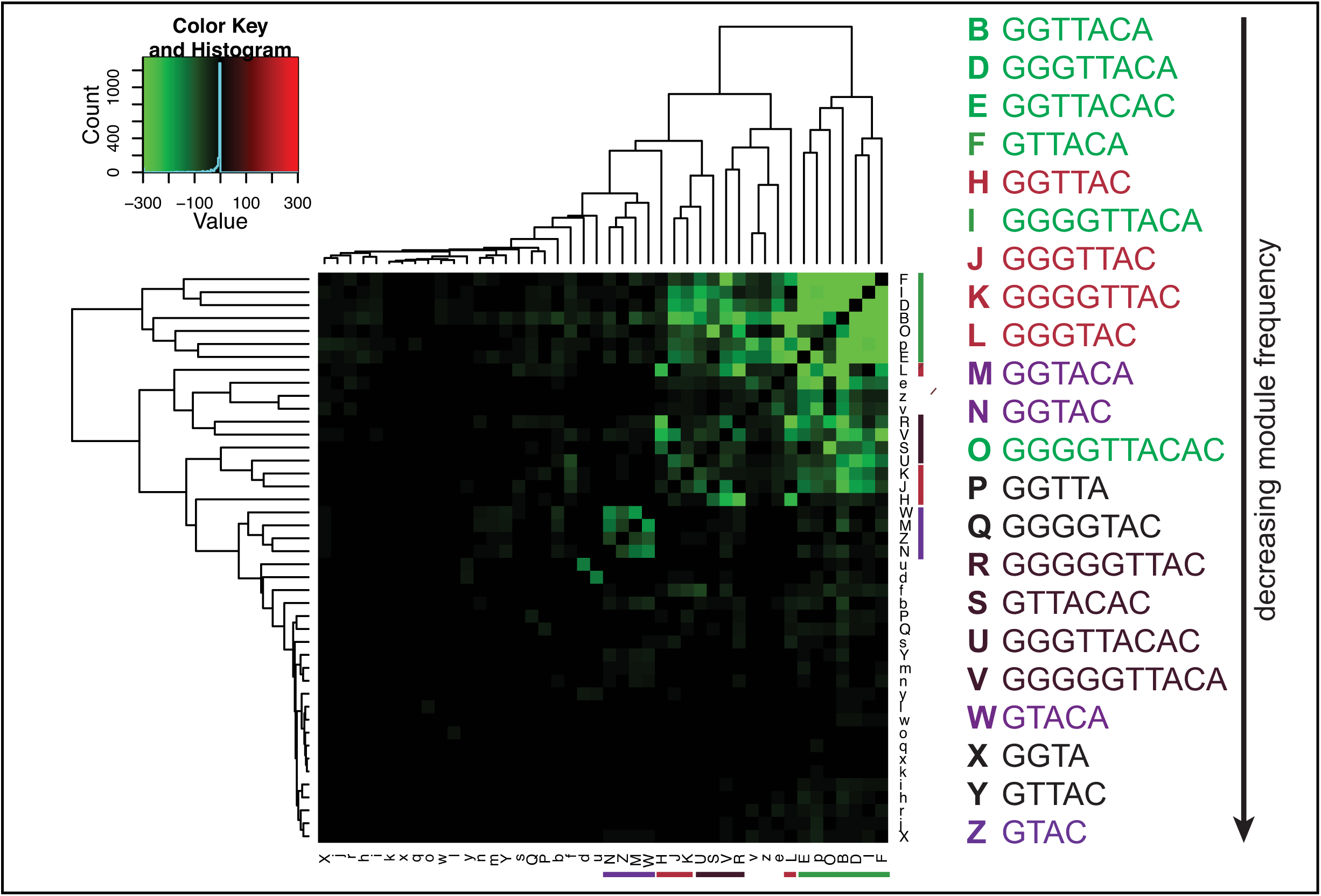
Association analysis reveals enrichment of “like” repeats within individual telomeres. The likelihood that each of the top 44 repeat modules coexist within individual telomere sequences was analyzed, hierarchically clustered and plotted according to the p-value of the association. Distinct families of repeat modules are color-coded: canonical repeats in green, AC modules in red, single T modules in purple and poly G and ACAC modules in brown (see text).

**Figure 4.**
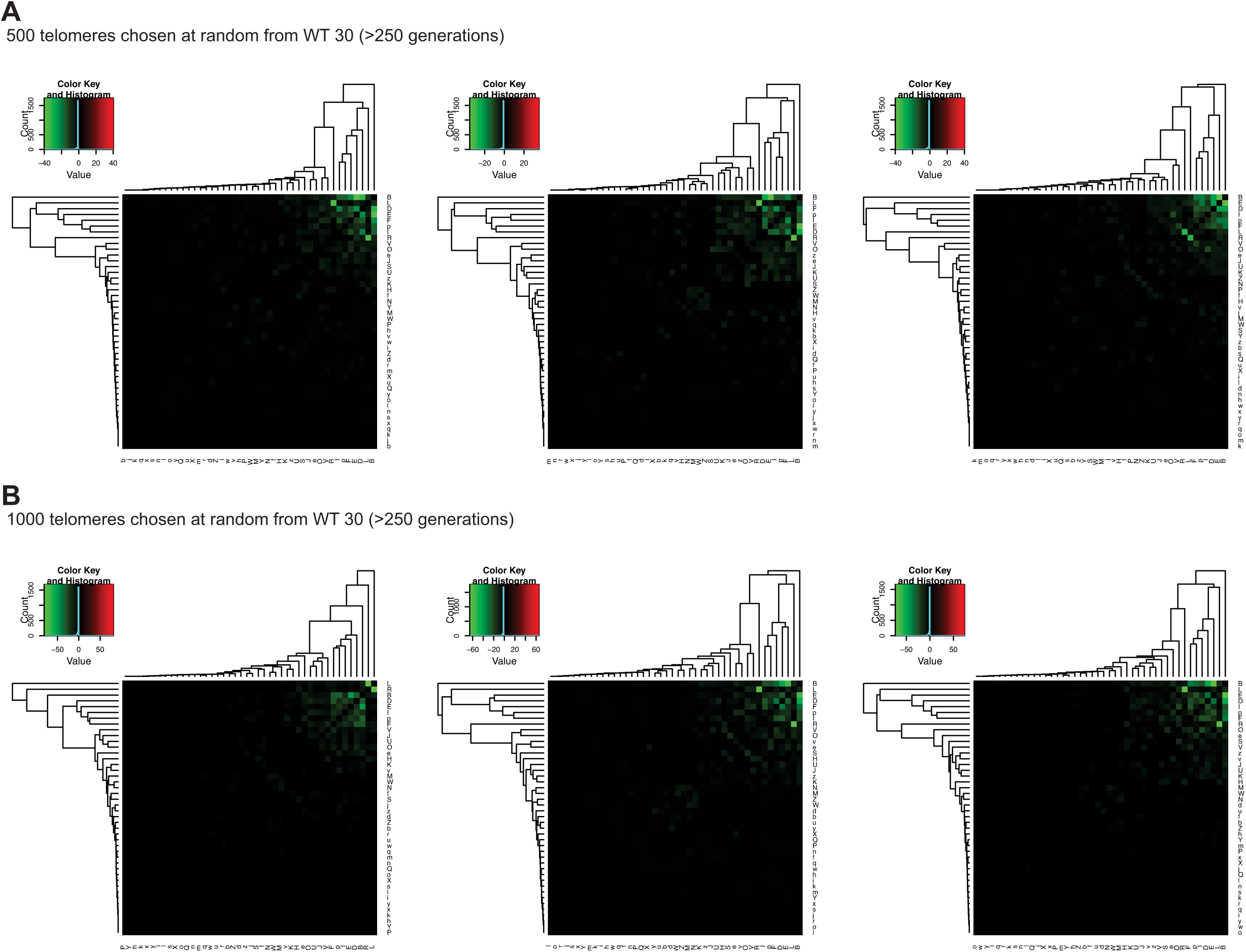
Robust clustering of modules requires thousands of telomere sequences. (A) Three representative examples of telomere association analysis from 500 sequences taking from the data plotted in Figure 3. (WT >250 generations) (B) Three representative examples of telomere association analysis from 1000 sequences taking from the data plotted in Figure 3. (WT >250 generations)

**Figure 5.**
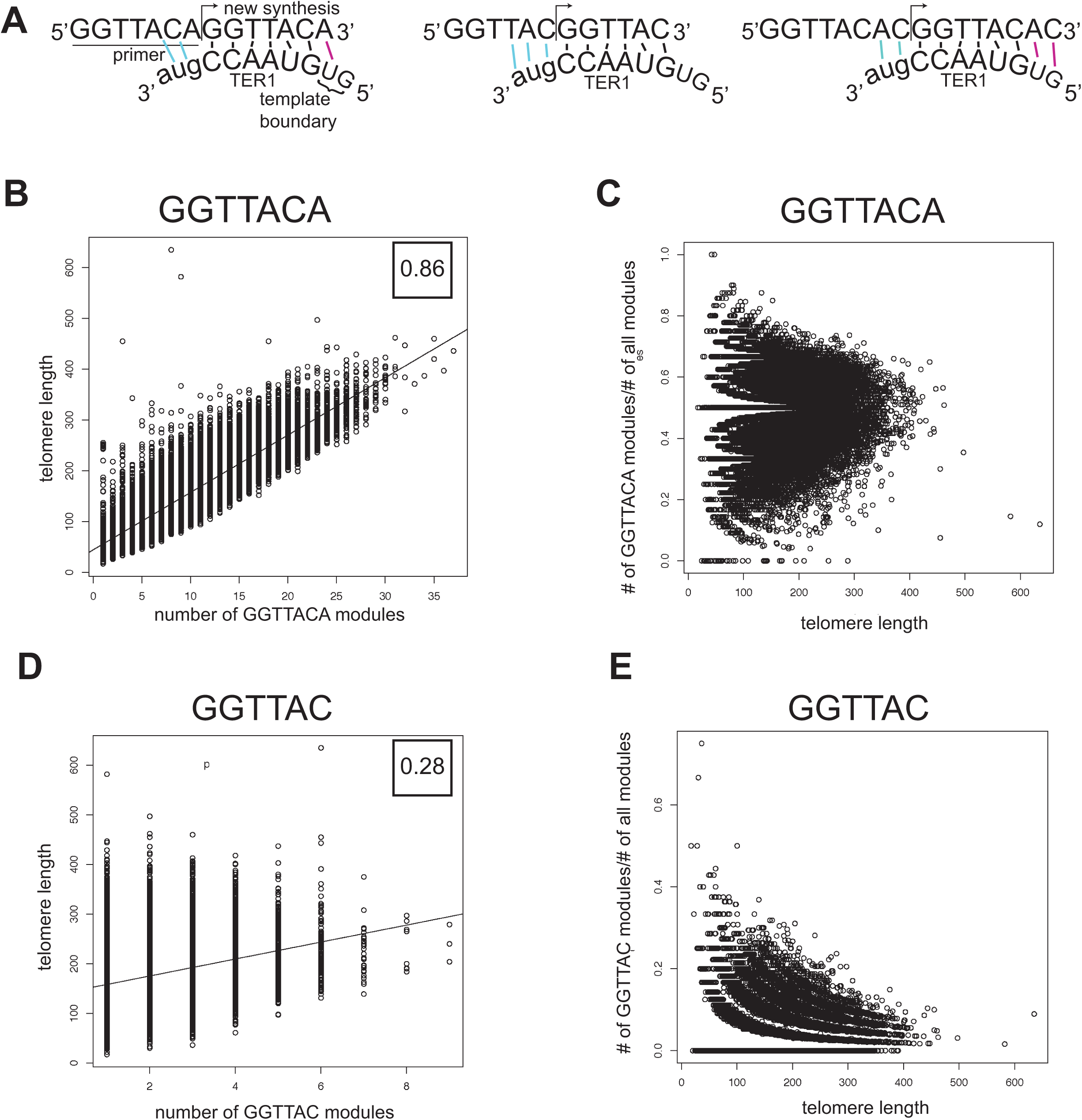
(A) Cartoon of the chromosome end (primer) aligned to the TER1 template RNA for three of the repeats: GGTTACA (left), GGTTAC (middle) and GGTTACAC (right). (B) Plot of telomere length versus the number of the GGTTACA repeat module per telomere. (C) Plot of the fraction of GGTTACA modules per telomere as a function of telomere length. (D) Plot of telomere length versus the number of the GGTTAC repeat module per telomere. For (B) and (D) the Pearson’s correlation value, which is a measure of the linear relationship between the number of the given repeat module and telomere length, and has a maximum value of 1, is included. (E) Plot of the fraction of GGTTAC modules per telomere as a function of telomere length.

**Figure 6.** Loss of telomerase activity influences telomere composition. (A) Representation of the telomere repeat content of *trt1Δ* telomeres shortly after germination (22 generations) plotted as in Figure 2A. Each circle is equivalent to 1%. (B) Plot of module composition in WT compared to *trt1Δ* telomeres (n=3 libraries for WT and n=2 libraries for *trt1Δ).* Plotted as the mean with standard deviation. Significance was determined by multiple t-tests with a false discovery rate of 5%. * p<0.05, ** p<0.01, **** p<0.0001 (C) Association analysis of the repeat modules of *trt1Δ* telomeres plotted as in Figure 3. Loss of telomerase activity leads to the emergence of a subpopulation of telomeres that are enriched for modules ending in ‐AC and – ACAC.

### Telomere PCR (Telo-PCR) and sample preparation

As shown in Figure 1, telomere ends were first ligated to an anchor primer (5’-[Phos]AGGGTTAATAAGCGGCCGCGTCG[AmC3]-3’) using CircLigase II ssDNA ligase (Epicentre) followed by PCR amplification and purification. Each 20 μl ligation reaction contained 10 μg of genomic DNA from phenol/chloroform extraction and 0.25 pM anchor primer and was executed as suggested by the manufacturer using the included buffers (Epicentre), including addition of betaine. The ligation reaction was incubated at 60°C for 2 hours, followed by 10 minutes at 80°C to terminate the reaction. Anchor-ligated telomeres were PCR amplified using conventional Taq polymerase (Roche; with 5% DMSO added), LongAmp Taq 2X Master Mix (New England Biolabs) or iProof (Bio-Rad; using “GC buffer”), with subtelomeric primer “1” (5’-CGAGGCTGCGGGTTACAAGGTTACG-3’) or “2” (5’-GGGAATTTAGGAAGTGCGGTAAGTTG-3’) and the reverse complement of the anchor sequence (5’-CGACGCGGCCGCTTATTAACCCT-3’), as described in Table 1 and below. In each 50 μl PCR reaction, 2 μl of ligation product was added. Different PCR cycling conditions were carried out depending on the polymerase. For Taq/primer “1”: 95°C (3 min), followed by 45 cycles of 95°C (30 sec), 52°C (30 sec) and 65°C (40 sec), and a final extension at 65°C (5 min). For iProof/primer “2”: 98°C (3 min), followed by 45 cycles of 98°C (10 sec), 60.5°C (15 sec) and 72°C (45 sec), and a final extension at 72°C (5 min). To prepare samples for high-throughput sequencing, 6 tubes of 50 μl PCR products were pooled together and concentrated and buffer exchanged using an Amicon Ultra 0.5 mL centrifugal filter (100,000 NMWL; Millipore) to remove free primers. The resulting eluate was diluted by addition of 100 μl of sterile water and further purified by Qiagen PCR purification kit.

**Table 1:**
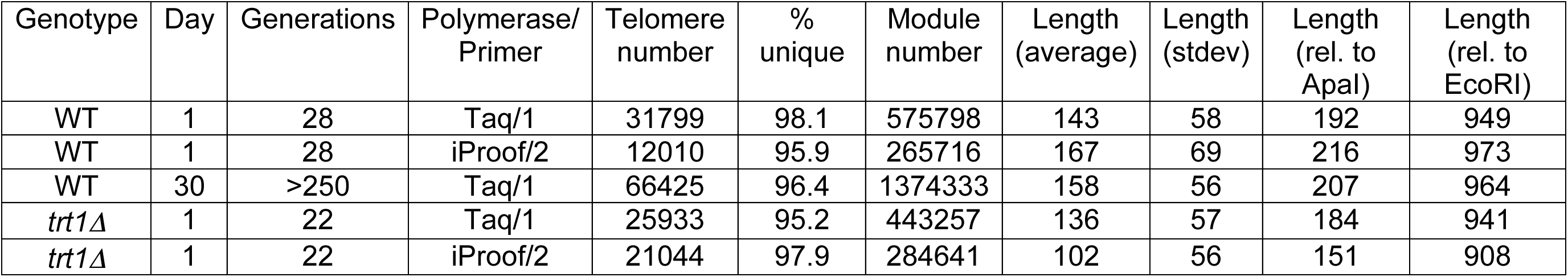
Telomere library information, length reported from start of telomeric repeat tract (see Supplemental Material)

| Genotype | Day | Generations | Polymerase/<br>Primer | Telomere<br>number | %<br>unique | Module<br>number | Length<br>(average) | Length<br>(stdev) | Length<br>(rel. to<br>ApaI) | Length<br>(rel. to<br>EcoRI) |
| --- | --- | --- | --- | --- | --- | --- | --- | --- | --- | --- |
| WT | 1 | 28 | Taq/1 | 31799 | 98.1 | 575798 | 143 | 58 | 192 | 949 |
| WT | 1 | 28 | iProof/2 | 12010 | 95.9 | 265716 | 167 | 69 | 216 | 973 |
| WT | 30 | >250 | Taq/1 | 66425 | 96.4 | 1374333 | 158 | 56 | 207 | 964 |
| *trt1Δ* | 1 | 22 | Taq/1 | 25933 | 95.2 | 443257 | 136 | 57 | 184 | 941 |
| *trt1Δ* | 1 | 22 | iProof/2 | 21044 | 97.9 | 284641 | 102 | 56 | 151 | 908 |

### PacBio sequencing and initial pipeline for analysis of telomere sequences

Purified telomere PCR products were used to generate libraries for high-throughput sequencing at the Yale Center for Genome Analysis. SMRT libraries were generated and sequenced according to the manufacturer (PacBio) using the protocol for ~250bp libraries at the Yale Center for Genome Analysis. The circular consensus sequence was determined using the manufacturer’s analysis pipeline to give raw telomere sequences. Sequences that did not contain the amplifying primers were excluded, with up to two mismatches allowed during the primer search. Sequences were further filtered based on the presence of at least one GGTTAC. The subtelomeric sequence upstream of (and including) the primer “1” sequence was trimmed off and the first base pair after this primer was assigned the value of “1 bp” for the purpose of measuring telomere length. The pipeline was written in Perl and plots were generated using R. For fitting of log normal plot (Fig. 1D) we used the equation: Y=Amplitude*exp(-0.5*(ln(X/Center)/Width)^2); the 95% confidence intervals were: amplitude: 488.4 to 497.2; center: 131.9-133.0; width: 0.3820-0.3899. All programs will be made freely available.

### Southern blot

Telo-PCR reactions were electrophoresed on a 1% agarose gel, denatured and transferred to Zeta-Probe GT membrane (Bio-Rad). The membrane was UV-crosslinked and hybridized with a telomere-DIG probe at 42°C for 2 hours using ExpressHyb Solution (Clontech). The probe was amplified by PCR in the presence of DIG-dUTP (DIG DNA Labeling and Detection Kit, Roche) from a template plasmid having 260 bps of telomeric repeats. After hybridization, the membrane was washed and incubated with anti-DIG-AP-conjugate (1:1000 dilution). The blot was then treated with SuperSignal West Femto Maximum Sensitivity Kit (ThermoFisher/Pierce), and imaged on VerSaDoc imaging system (BioRad) using the software Quantity One 4.6.9.

### Telomere module detection and analysis

Modules were detected using a script written in Perl (available upon request). The junctions between modules were assigned based on G motifs (G_1_→_n_), with new modules occurring at the first G after the previously assigned module, moving in a 5’ to 3’ direction. All detected modules were ranked according to the frequency of occurrence in each library. The coverage of each module was calculated by its occurrence multiplied by its size, and divided by the total nucleotides in the sample.

### Association analysis

The top 44 modules from the WT sample (>250 generations) were selected for association analysis. For each individual telomere sequence, presence/absence of the top 44 modules was evaluated in a binary fashion and the Fisher’s test was performed to derive a p-value for each pairwise combination. The association relationships were subjected to hierarchal clustering analysis and are displayed as a heat-map with a scale based on the p-values.

## Results

### Next-generation sequencing and analysis of telomeric repeats

To generate large datasets of the telomeric repeat content of individual *S. pombe* telomeres in their entirety, we developed a method that adapts established telomere cloning protocols (19,21) for next-generation sequencing (“TeloPCR-Seq”; Figure 1A). First, we carried out ligation-anchored PCR on genomic DNA templates using subtelomeric primers designed based on a cloned reference sequence (24) (Figure 1A and Supplemental Material). Products amplified using a subtelomeric primer just upstream of the telomeric repeat tract have a range of sizes (Figure 1B). We validated that these products are both dependent on the anchor primer and can be hybridized to a telomeric probe by Southern blot (Figure 1C). The amplified telomere products were then used to create circular libraries and sequenced using the singlemolecule PacBio RS platform.

We first analyzed a telomere library generated from wild-type (WT) cells that had been cultured for over 250 generations (Table 1). The raw dataset contained more than 60,000 individual sequences that contain at least one telomeric repeat (defined as the minimal GGTTAC sequence); the vast majority of these sequences were assigned as unique (96.4%, Table 1). The average length of all telomeres was 158 +/‐ 56 base pairs (bps) from the end of primer “1” (Figure 1D, cyan, equivalent to 207 bps from the Apal site and 964 bps from the EcoRI site in the subtelomere). Plotting the observed frequency of a given telomere sequence versus its length revealed a distribution best described with a log-normal model (Figure 1D, red line; R square = 0.988), with a degree of skew towards longer telomeres. The small deviation from the mean and nearly normal distribution of the data are consistent with observations that telomerase activity is tightly regulated to achieve telomere length homeostasis (25). This deep data set reveals the presence of both critically short and very long telomeres within a normal, WT population.

We wanted to investigate whether our initial analysis was missing sequences from shorter telomeres that could potentially lack the subtelomeric “1” priming site (see also Supplemental Material). Comparing the sequences derived from products produced using primer “1” with those amplified using the second, upstream subtelomeric primer (“2”; Figure 1D, green, and Supplemental Material), we found the average raw telomere length to be 247 +/‐ 69 bps, equivalent to an average length of 157 bps from primer “1”, nearly identical to the predicted size of 158 bps based on the known subtelomere sequence (24) (Table 1 and Supplemental Material). Thus, primer “1” seems to capture the vast majority of telomere sequences from WT cultures, suggesting that telomeres rarely loose their entire telomeric repeat track due to replication and/or nucleolytic degradation. In addition, analysis of the subtelomeric sequence between primers “1” and “2” provided the opportunity to assess the quality of the sequencing data. Analysis of over 12,000 sequences derived from the primer “2” data set revealed that the “in between” subtelomeric region was identical to the previously reported sequence (24) for 82% of sequence reads and was greater than 90% similar for over 98% of the sequence reads. It is highly possible that the imperfectly matched sequences represent individual telomeres that vary from the cloned database reference sequence, as these products are expected to be amplified from both chromosome ends of chromosomes I and II, whereas the cloned telomere sequence arose from a single telomere. However, we cannot rule out that sequencing error contributes as well.

### The telomeric repeats are composed of a small number of module variants

To facilitate analysis of the telomeric repeats, we developed a routine in Perl that breaks down each individual telomere sequence into its component repeat units. Essentially, after trimming off of the primer sequences, the first telomeric repeat variant or “module” at the 5’ end was assigned. Additional modules were assigned from the 5’ to 3’ end. Using this method, we assessed repeat preference for individual repeat modules from libraries prepared either shortly after germination (28 generations) using either Taq polymerase and primer “1” or iProof polymerase and primer “2” (followed by removal of the subtelomeric sequence), or after extensive culturing of cells (250 generations); the latter library contained over 660,000 variant modules (Table 1). Consistent with previous studies (13,16) and the sequence of the TER1 RNA template region, the most common module is GGTTACA, which makes up nearly half of all repeats in WT cells (Figure 2A and B); combined with the G number variant GGGTTACA, these two repeats cover about 60 percent of the total sequence. In all, the top eight repeat modules cover about 85% of the total modules and vary in length from 6-9 nucleotides. As the 3’ end of the module reflects the use of the TER1 template boundary (10-12), we analyzed the prevalence of modules ending in ACA, AC, or ACAC. In all libraries, modules ending in ACA made up about 75% of the total, with AC endings (excluding ACAC endings) making up ~10% and ACAC endings making up only ~5% (Figure 2C). In general, this global analysis of telomere content is very similar to that obtained by traditional telomere cloning and Sanger sequencing (13,18). Plotting the distribution of module length reveals that about ~85% of fission yeast repeats are between 6-8 nucleotides in length (Figure 2D). This variability in repeat length and sequence is more regular than that seen in *S. cerevisiae* (although greater than that seen in mammalian systems), perhaps reflecting the greater similarity between fission yeast and mammalian telomere binding proteins such as Pot1 and TRF1/2, which are absent in budding yeast (26). Importantly, module usage was highly similar regardless of the primer and polymerase employed for telomere amplification (Figure 2A, compare WT (t) and WT (i) samples). We next wanted to investigate how sequencing error may influence the accuracy of module assignment. If we assume that sequencing error is random with respect to nucleotide and position (27), we can predict all of the modules that could occur by a single base pair error in the most common GGTTACA module (for example, GCTTACA). If we consider all modules that are equivalent or are a one base pair change from GGTTACA, we find that GGTTACA makes of over 96% of all such modules (Supplemental Table 1). Based on incompatibility with the TER1 template, we assume that all single base pair errors within the GGTTACA module would occur by sequencing error rather than due to biological variability (although it remains possible that these modules arise due to telomerase). The most prevalent of these, GG<u>G</u>TACA, makes up <0.4% of all modules (Supplemental Table 2). Thus, modules that comprise less than 0.4% of the module population could be accounted for by sequencing and/or analysis error (the top 23 modules are above this threshold), including all modules that we interpret to be biologically important (Supplemental Tables 1 and 2, and see below).

Consistent with the uniqueness within the libraries, individual telomeres were not readily assigned to families with substantial tracts of highly conserved sequence. However, it is also possible that we overestimate uniqueness due to errors that occur during amplification, sequencing, or consensus alignment. To mitigate these concerns, we chose to take further advantage of the power in analyzing module distribution within entire individual telomeres. Consistent with true biological variability in the telomere population, while the first telomeric repeat downstream of the “primer 1” sequence is common for nearly 90% of telomeres (the GGTTACAC module), the identity between telomeres rapidly falls off with increasing repeat number (Table 2), similar to previous observations in budding yeast, in which centromere-proximal sequences remained constant while centromere-distal regions were subject to sequence change (22). This trend appears exaggerated in fission yeast telomeres, though this could also be attributable to methodological differences, as the budding yeast sequences were amplified from a single telomere rather than the mixed telomere population amplified in our approach (21). In addition, most repeat modules appeared to be uniformly distributed across the telomeric repeat tract without bias for the 5’ or 3’ ends, with the exceptions of GGTTACAC (which is biased to the 5’ end due to its prevalence as the first module) and the GGGTTACA and GTTACA modules (which appear biased internally within the repeat tract) (Figure 2E).

**Table 2:** Identity of module at positions 1-6 after subtelomere Highlighted values represent instances in which >50% of the indicated module is present at that site across the population

| Module number | GGTTACA | GGGTTACA | GGTTACAC | GTTACA | GGTTAC | GGGGTTACA | GGGTTAC | GGGGTAC | other |
| --- | --- | --- | --- | --- | --- | --- | --- | --- | --- |
| 1 | 2.0 | 0.2 | 89.1 | 0.4 | 0.8 | 0.2 | 0.2 | 0.2 | 6.9 |
| 2 | 90.5 | 0.3 | 0.4 | 1.9 | 0.8 | 0.3 | 0.2 | 0.2 | 5.4 |
| 3 | 88.2 | 0.9 | 0.2 | 1.8 | 2.8 | 0.2 | 0.2 | 0.2 | 5.5 |
| 4 | 84.3 | 3.9 | 0.2 | 1.9 | 1.8 | 0.2 | 0.6 | 0.6 | 6.2 |
| 5 | 31.3 | 43.2 | 0.6 | 1.6 | 0.8 | 0.7 | 0.6 | 0.6 | 19.6 |
| 6 | 54.7 | 22.9 | 0.6 | 5.8 | 1.2 | 0.9 | 1.2 | 1.2 | 11.1 |

### “Like” modules are more likely to present within individual telomeres

Next, because this approach reveals the repeat sequences from individual telomeres in their entirety, we wished to explore whether repeat module usage at the single telomere level could be explained entirely by their relative prevalence in the population. To facilitate this, we analyzed the probability that two modules were present in the same telomere pairwise, giving rise to a value representing the p-value of each association. This matrix of values was then subjected to hierarchal clustering and scaled to reveal the most highly associated modules (Figure 3). The top 44 modules were given alphabetical identifiers (the letters A, C, G, and T were excluded, upper case followed by lower case), as displayed, in the same order as their overall abundance (Supplemental Table 2). As would be expected, the most prevalent modules clustered together and had a high significance of association (for example, the top four canonical modules B, D, E and F and less common I and O, colored green).

Interestingly, other “families” of modules could not be explained simply by their abundance. Most obviously, the fifth most prevalent module, GGTTAC (module “H”) was not associated with the other top modules, instead having more significant association with less common modules ending in AC (J, K and L, colored red). Thus, modules ending in AC are more likely to occur in telomeres with other 3’ AC modules and less likely to present with the more common 3’ ACA modules. The observation of “like” modules occurring most prevalently together extends beyond this example; indeed, the rare modules containing one T (M, N, W and Z, colored purple) also had significant association with each other. A fourth family was enriched in modules ending in ACAC and extended G-stretches (R, S, U and V, colored brown). As predicted from this analysis, we observe that rare modules (such as the u and d modules that include three Ts) can be found highly enriched in a small number of telomeres far above their prevalence in the total population (see Supplemental Material).

We next assessed how the information revealed by high-throughput telomere data sets compared with results that could by obtained by conventional sequencing. Taking either 500 or 1000 telomeres sequences chosen at random from the full data set analyzed in Figure 3, we found that, although the most abundant modules showed high association (as would be predicted), the ability to discern association between “AC” ending modules and other more rare modules was lost (Figure 4A and B). Thus, the full value of the association analysis of module families requires the depth of sequencing afforded by this method.

### The GGTTACA but not GGTTAC module positively correlates with length

We were interested to explore the unusually poor correlation between the “H” module, GGTTAC, and the other canonical modules that end in ACA. Modules ending in the AC ending rather than the more common ACA ending would be predicted to increase the affinity of chromosome end for the TER1 RNA, as they provide the most optimal base pairing with the AUC of TER1 (cyan lines, Figure 5A), potentially increasing the efficiency of telomerase elongation and/or processivity. When we analyzed the number of GGTTACA modules per telomere, we found that prevalence of GGTTACA is well predicted simply as a function of length (Figure 5B, Pearson’s correlation of 0.86). If we analyze the fraction of GGTTACA modules per telomere as a function of length (Figure 5C), we see a distribution that starts out with a high variance at very short telomeres (likely due to a small number of total modules) but converges around a horizontal line at a fraction of around 0.5, the fraction that is observed in the total population (Figure 2A). Interestingly, when we analyzed the prevalence of the GGTTAC module as a function of length, we found that the two were poorly correlated (Figure 5D, Pearson’s correlation of 0.28). Further, the fraction of telomere modules made up of GGTTAC is highest in short telomeres and decreases with increasing telomere length (Figure 5E). These differences may reflect preferences for repeat module addition by telomerase, perhaps based on TER1 primer annealing, that are influenced by the telomere state. Alternatively, the AC ending could antagonize sequence degradation from the 3’ end or stabilize end structure, in which case this module could be enriched by loss of the extreme 3’ A from ACA endings.

### Loss of telomerase activity rapidly leads to altered telomere sequences

To test the ability of this telomere sequencing approach to contribute to our understanding of telomere biology, we next assessed how loss of telomerase activity influenced telomere sequence. To this end, we sequenced the telomeres of cells lacking the catalytic component of telomerase, Trt1, shortly after germination (Figure 6). Investigating the acute changes in telomere sequence was necessary, as most *S. pombe* cells lacking Trt1 ultimately survive through chromosome circularization; although maintenance of telomeres by recombination also occurs, this mechanism of survival is generally not stable in the absence of additional genetic perturbations (28,29). Thus, we examined the composition of *trt1Δ* telomeres early after spore germination (22 generations) before chromosome circularization ensues to allow us to investigate telomere length and module usage as cells enter the growth crisis; we expect that in this phase telomeres are shortening due to both the “end-replication problem” as well as nucleolytic degradation. Even after these relatively few cell divisions, the mean *trt1Δ* telomere tract length is shorter than that seen for WT cells (136 +/‐ 57 bps using “primer 1” and Taq or 102 +/‐ 56 bps using “primer 2” and iProof; Table 1). Interestingly, we also see changes in module usage, with a decrease in the canonical GGTTACA and GGGTTACA (ACA ending) repeats and an increase in the GGTTACAC, GGTTAC and GGGTTAC (AC ending) repeats (Figure 6A and B). While the prevalence of GGTTACAC can be explained by telomere shortening and the enrichment of this module at the far 5’ end of the telomeric repeats (Figure 2E), the prevalence of GGTTAC cannot. When we subjected the *trt1Δ* telomeres to module association analysis followed by hierarchical clustering, we observe that the GGTTAC containing telomeres make up a unique family that cluster distinctly away from telomeres rich in “canonical” repeats (Figure 6C). Because these cells lack telomerase activity, this observation supports the notion that the enrichment of modules ending in AC reflects either an increased resistance to nucleolytic degradation and/or arises as a consequence of telomere recombination.

## Discussion

Next generation sequencing is an extraordinarily powerful method to address genome biology. However, the inability to assign and assemble repetitive sequences back to their starting structure remains a major hurdle for applying such platforms to the study of repetitive elements. Although analysis of short telomeric repeat reads has revealed shifts in telomere variant sequences between cells that maintain their telomeres through telomerase or the alternative lengthening of telomeres (ALT) pathway (30,31), suggesting that changes in telomere content is biologically meaningful, this method can neither support assembly of these sequences back to the initial telomere sequence, nor can it exclude possible contributions from internal telomeric repeat tracts.

Here, we have developed methods to both sequence and analyze the telomeric repeats at the level of individual telomeres in the model organism, *S. pombe.* Simple modifications to this approach should make this method useful for other model systems. The experimental requirements for application to other models include having knowledge of subtelomeric sequence and the ability to identify a suitable, unique priming site. The primer may be specific to a single telomere or present in multiple telomeres (as is the case here); both situations can be experimentally valuable. In addition, because the TER1 template sequence is highly variable among yeasts (and distinct from mammalian systems), adjustments to the computational routines that decompose telomere sequences into their repeat modules will need to be made. Further, telomere length must be considered. Based on the PacBio technology, we anticipate that telomere repeat tracts up to ~ 5 kilobases from the subtelomeric primer should be possible, although sequencing accuracy will decrease because large inserts will have lower coverage in the sequential sequencing of the circular template that is essential to improve accuracy over the single pass PacBio error rate of ~15% (32). Even so, because the errors in PacBio are random and can be accounted for (27), and because this method can uniquely provide long-read information, it should nonetheless prove a powerful tool for examining telomere repeat content in many organisms.

This high-content telomere sequencing approach reveals an exquisite control over telomere length across a cell population (Fig. 1D), while at the sequence level individual telomere content is in fact highly diverse (Table 1). Despite this sequence complexity, our analysis suggests that there is a strong association of “like” modules within individual WT telomeres (Fig. 3). While the underlying mechanisms are not yet clear, we propose three plausible (and not mutually exclusive) models. First, the processive nature of telomerase (33,34) raises the possibility that distinct forms of telomerase (perhaps through different TER1 RNA conformations) will add multiple repeats of a given type, leading to enrichment of that repeat type in individual telomeres. Consistent with this possibility, we see that the same rare modules are sometimes added in tandem (see Supplemental Material). Second, perhaps the existing telomere sequence promotes recruitment of telomerase conformations that recognize and reiterate the repeat type, making it more likely that the same repeat type is added. Third, telomere-associated factors such as the multi-subunit shelterin complex (9) may promote recruitment of specific telomerase conformations. Further *in vitro* experiments to evaluate whether the repeat modules added by *S. pombe* telomerase are influenced by the initial primer sequence and/or the presence of shelterin components will be required to address these questions.

One open question is whether module composition is biologically important, particularly the enrichment of 3’ AC containing modules seen in short telomeres (Figure 5). If the terminal telomeric DNA repeat (or the “primer” *in vitro)* ends in AC or ACAC, this could enhance base pairing with the template RNA (Figure 5A). Such an increase in affinity between the newly synthesized terminal repeat and the TER1 RNA could, in theory, promote the efficiency of telomerase-mediated elongation and/or telomerase processivity. Alternatively, a chromosome end that terminates in an AC or ACAC ending could promote enhanced recruitment of telomerase. Indeed, this latter possibility is intriguing because such a mechanism could contribute to the observation that telomerase recruitment is biased towards the shortest telomeres (35). Importantly, there is evidence for a correlation between the 3’ end of a given module sequence (AC/ACAC versus ACA) and the extent of G-stutter in the following repeat module, with G-stuttering more commonly found after ACA endings (18), suggesting that “primer”-TER1 base pairing can influence the subsequent module added by telomerase. It is important to note, however, that targeted disruption of base pairing between the DNA primer and template RNA in budding yeast telomerase negatively influences telomerase elongation *in vitro,* but does not disrupt processivity (36). This mirrors observations for purified telomerase from other sources *in vitro,* which suggest that the affinity of telomerase for oligonucleotide substrates is largely unaffected by the extent of base complementarity between the primer end and the TER1 RNA (37,38). As the possibility that the primer sequence influences elongation and processivity has not yet been thoroughly investigated in fission yeast *(in vitro* or *in vivo),* particularly for these specific primer sequences (GGTTAC/GGTTACAC versus GGTTACA), testing this interpretation awaits further experimentation. Importantly, our finding that modules ending in AC are more prevalent in shorter telomeres in both WT cells and those lacking telomerase activity suggests that, alternatively or in addition, these modules may arise from degradation of the 3’ A rather than differential addition by telomerase. Again, this may highlight a difference with budding yeast, as a previous study in this model system (in which telomerase promotes incorporation of degenerate TG_2-3_(TG)_1-6_ repeats) found that G or T were found equally at the extreme 3’ base position in both the presence or absence of telomerase activity (21). We expect that the method described here for high-content analysis of telomere sequences from fission yeast, combined with further *in vitro* experiments, will together prove invaluable for examining the mechanisms and consequences of repeat variability in *S. pombe.*

## Accession Numbers

Raw sequence data is available at the NIH SRA database, accession # SRP043421.

## Acknowledgements

We are indebted to the Yeast Genomic Resource Center (YGRC) at Osaka University for providing access to strains, as well as the many researchers who have deposited strains at this resource. We also thank Dr. Claus Azzalin and Dr. Patrick Lusk for valuable feedback on the manuscript.

